# Adaptive therapy in cancer: the role of restrictions in the accumulation of mutations

**DOI:** 10.1101/2023.05.18.541330

**Authors:** David Fontaneda, Ramon Diaz-Uriarte

## Abstract

**BACKGROUND:** Cancer is currently one of the leading causes of premature death in the world, and is predicted to continue rising even despite the continuous discovery of novel treatments. New approaches, like adaptive therapy, try to minimize the problem of drug resistance, but there are still many open questions and unstudied phenomena that need to be tackled in order to make this approaches viable in real patients; among these, the possible effects that restrictions in the order of accumulation of mutations could have.

**RESULTS:** We have developed a spatially explicit agent-based model capable of simulating tumor growth and adaptive therapy in a highly flexible way. We show that when we consider restrictions in the order of accumulation of mutations and their effect in tumor architecture, the predicted genotypes of the cells that are inhibiting the growth of resistant cells can be very different to the ones predicted by perfectly mixed models.

**CONCLUSION:** We identify a divergence between the expected and real genotypes of the cells inhibiting the growth of the resistant population that has not been previously documented. This effect, if not taken into account, could negatively affect our predictions of adaptive therapy success and could hinder our advances in the development of new approaches to improve adaptive therapy. This discovery suggests the need for more studies that take into account the spatial component of cancer, specially when dealing with tumors with high heterogeneity. Furthermore, our model is able to simulate scenarios of tumor development and adaptive therapy, making it useful both for research and for education.

## 1 Introduction

The term cancer refers to an ample group of diseases in which cells divide and grow uncontrollably due to accumulative genomic and epigenomic alterations [17]. It is estimated that almost 10 million annual deaths can be attributed to these diseases, making it the leading cause of premature mortality in most developed countries [42]. Moreover, this number is predicted to continue growing, reflecting both an increase in life expectancy worldwide along with a decrease in the other leading causes of death such as stroke and coronary heart disease [7].

The preferred strategy for cancer treatment varies depending both on the specific tumor type and the specific situation of the patient, such as age or cancer stage [31]. The most common modalities can be classified as either surgery, radiation therapy or systemic treatment, this last one including chemotherapy, targeted therapy, hormonal therapy and immunotherapy [31]. Although systemic treatments are not available for all cancer types, and often have harsh side effects on the patients, they have led to an improvement in overall survival, specially in liquid and metastatic tumors, where the effective use of other alternatives becomes more complicated [35]. One vital problem, however, is that during their evolutionary process, cancerous cells frequently end up developing resistance to the treatment, leading to the loss of control of the population and, ultimately, the death of the patient [27]. There have been many proposals to tackle this problem and one of the most promising ones is “adaptive therapy” [19].

“Adaptive therapy” is an approach inspired by other fields with similar problems, like antibiotic resistance and agricultural pest control [19, 22, 23, 43]. Adaptive therapy shines in scenarios where the emergence of drug-resistant cell populations is almost certain, and in consequence, achieving complete eradication of the population is improbable. In these cases, we shift the goal from curing the patient to prolonging it’s quality life span [22]. To achieve this, adaptive therapy exploits the competition between the different cell populations coexisting in the tumor, deliberately stopping the treatment early and maintaining an “acceptable burden” tumor size. The competition for resources slows down the growth of the resistant population and is able to effectively delay the progression of the tumor, defined as the moment we are no longer able to keep the tumor under control [40].

Historically, due to the complexity of studying cancer as an evolutionary disease, most of the insight into this approach has been gained thanks to mathematical models and computational simulations [3, 18, 19, 40, 41, 45, 48]. However, in recent years adaptive therapy has also been shown to produce an increase in time to progression (TTP) in preclinical studies for breast and ovarian cancer [15, 19], and there are multiple ongoing and planned treatment trials to test this approach [48] in rhabdomyosarcoma (NCT04388839), castration-sensitive prostate cancer (NCT03511196), BRAF mutant melanoma (NCT03543969) and ovarian cancer (NCT05080556), showing an increasing interest in this approach among the cancer research community.

The studies cited above do not consider possible restrictions in the order in which mutations accumulate in cancer. The multi-step nature of tumorogenesis has been a long known fact [1]; it is also known that the order in which these steps take place is not random. Instead, the appearance of driver mutations is restricted in such a way that only some sequential orders are allowed, and the appearance of certain mutations in the population makes other mutations much more likely [4, 16].

The identification of these restrictions is an active field of study. Cancer progression models (CPMs) have been developed to try to infer these restrictions by analyzing genotype frequency data from cross-sectional samples of tumor biopsy, a common data source available for a lot of cancer types; see [4, 5, 13] for overviews. These models open the door for prediction of tumor progression paths [10, 14, 28] and, among them, the prediction of paths leading to resistance-conferring mutations [21, 32]. These features make CPMs a specially interesting tool for the study of adaptive therapy. More immediately, restrictions in the order of accumulation of mutation could lead to different evolutionary dynamics that strongly affect the competitive patterns inside tumors.

The objective of this manuscript is to start examining how restrictions in the order of accumulation of mutations could affect adaptive therapy, and how CPMs can be leveraged to design more effective adaptive therapies. Here we focus specifically on the effect that these restrictions in the order of accumulation of mutations have on the divergence between the expected and real genotypes of the cells inhibiting the growth of the resistant population. To examine this problem, we use a flexible agent-based model (ABM) that can be used to address this question, as well as other questions that pertain to the use of adaptive therapy with explicit consideration of the spatial dimension.

## 2 Materials and methods

### 2.1 Agent-Based Modelling

To study adaptive therapy theoretically, we need to resort to models that are able to capture the properties of real tumors. The choice of the model depends on the specific part of the phenomena that one wants to study but, historically, the most common ones have been Lotka-Volterra equations [40], cellular automata [3] and agent-based models (ABMs) [18, 41].

Agent-based modeling [26] in particular is a framework where we model each cell as a separate individual, referred to as an “agent”. These agents occupy a specific place in space, possess individual properties and follow certain stochastic rules in response to their surroundings. In tumor models, this translates to each cell being able to have it’s own genotype, fitness and position in space, and being able to interact with other cells to decide whether it should migrate, reproduce, or die. This is particularly useful, as it allows us to explore the complex emergent properties that appear in real tumors without having to explicitly define them.

Agent-based models, however, also have their drawbacks. One of the main disadvantages of ABMs is that they are often more computationally intensive than other common alternatives, as we need to keep track of each cell individually. Also, because of their stochastic nature, small changes in the starting conditions can result in vastly different results. This means that, in practice, we need to average the results over a large number of simulations if we want to draw meaningful conclusions. Both of these effects combined make access to powerful hardware and a carefully optimized simulation software indispensable requirements to reduce the required simulation time to an acceptable timeframe.

### 2.2 General overview of our agent-based model

After careful review of the literature, we have not found any model that has studied the effect of restrictions in the order of accumulation of mutations on adaptive therapy [3, 18, 41, 45, 48]. Furthermore, recent work has shown that the spatial architecture of the tumor might play a bigger role in adaptive therapy than previously anticipated [3, 18, 41]. If we also want to study this effect we are restricted to only using models that explicitly model space, like cellular automata or ABMs.

With the objective of studying the potential effect of spatial architecture and restrictions in the order of accumulation of mutations on adaptive therapy, we have developed a new ABM that is able to implement all of these features in a flexible way. All the code has been written in Julia, a high-level programming language specialized in numerical analysis and computational science [6, 36]. For the development of the model, the library “Agents.jl” [9] has been extensively used. This library, a well tested collection of community-written code for designing ABMs, allowed us to implement our model in a robust way, reducing the chance of errors in the code while maintaining a very high flexibility compared with other alternatives.

In Figure 1 we show a “black box” model of our ABM, a representation that allows us to show the inputs and outputs of the model without considering the specifics of its inner workings, which we will explain later.

**Fig. 1.**
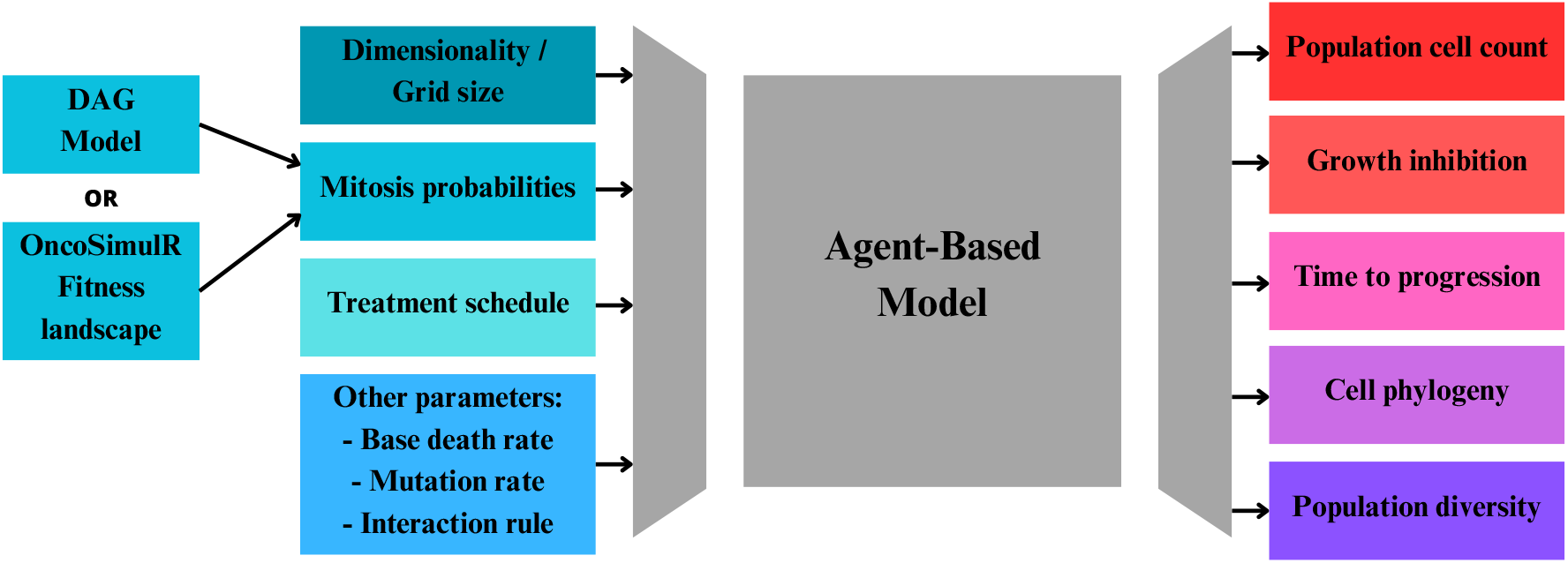
Black box model representing the functionality of our ABM. All the inputs of the model are shown on the left, and all data produced is shown on the right.

The most important inputs for our model are as follows:

- **Dimensionality and grid size**. With our model we are able to run simulations in 1D, 2D and 3D in arbitrarily large grids. In our model, each cell is able to occupy exactly one position of the grid. Therefore, this parameter implicitly limits the maximum allowed population.
- **Mitosis probabilities**. A table relating the genotype of a cell with its probability of starting mitosis. This table can be obtained either from the directed acyclic graph (DAG) that specifies the restrictions in the order of accumulation of mutations or directly from a fitness landscape such as the ones generated by the software package OncoSimulR [11].
- **Treatment schedule**. The specific details of the treatment, including whether we are using traditional continuous therapy, adaptive therapy, and the size we consider to be an “acceptable burden” for the patient.
- **Other parameters**. These include the base death rate of the cells, the migration rate and the mutation rate. We can also specify the interaction rules the cells will follow, such as contact inhibition [25], which mimics a scenario with strong replication inhibition due to spatial constrains, or the hierarchical voter model [37], representing a scenario where cells with driver mutations are unregulated and can proliferate at the cost of other cells with lower fitness.

Once the simulation of the tumor has ended, our model outputs data such as:

- **Cell count by genotype in each time step**. We can use this data to analyse the predominant population in each step and monitor the dynamics of the whole tumor size.
- **Resistant cells growth inhibition by genotype**. We can analyse the genotype of the cells that are inhibiting the growth of the resistant cells in each step.
- **Time to progression of the tumor**. Defined as done previously in similar models [18, 40], defining ”progression” as reaching an upper limit over the ”acceptable” size, which we take as a proxy for our inability to control tumor growth.
- **Cell phylogeny**. Allowing us to reconstruct the phylogenetic tree of all the cells that are present at the end of the simulations, and with them the true mutational paths that the population traversed to reach the treatment-resistant state.
- **Population diversity**. This provides statistics commonly used in ecology indicative of the diversity of genotypes and heterogeneity of the tumor, such as the species richness and the Shannon index at each step of the simulation.

All of this data allows us to analyse what parameters are essential for adaptive therapy success, and facilitate the search to find new, non studied phenomena that can be used to improve this approach.

### 2.3 Agent-based model (ABM) step loop

Our ABM has been implemented as a time-discrete model. In each time step, the complete list of agents is traversed, and the actions taken by each cell are decided according to the algorithm shown in Figure 2. The evaluation process taken for each cell in each time step can be described as follows:

1. First, each cell has a chance to be affected by random apoptosis [40] with a user given base death rate probability. If the cell is determined to be affected, it is instantly removed from the simulation. Otherwise, the algorithm continues.
2. Then, we check if there is space available around the evaluated cell according to the interaction rule specified. If we are using the contact inhibition model [25], we consider space to be available if there is at least an empty space in the Von Neumann neighbourhood [44] of the cell. If we are using the hierarchical voter model [37], a cell is able to colonise a space already occupied by other cell as long as this other cell has a lower fitness, thus, we consider neighbouring spaces with inferior cells to also be available. If there is no empty space, we end the evaluation for this cell, but if there is space available the cell has a chance to start a mitosis cycle.
3. We continue by using the same procedure as for random apoptosis to decide if the cell should reproduce. We generate a random number, and if is below the mitosis probability of the cell’s specific genotype, defined by the input fitness landscape, a mitosis cycle starts.
4. While mitosis is taking place, if the tumor is currently undergoing treatment and the cell does not have the resistance-conferring mutation, another random number is sampled to determine if the treatment affects the cell attempting division [41]. If the sampled number is below the kill rate of the treatment, the cell is instantly removed.
5. If the cell is still alive, the mitosis cycle continues. DNA is duplicated and there is a chance for each of the daughter cells to acquire random mutations according to the specified mutation probability [38]. Physical division is then completed and both daughter cells continue developing as separate agents. At this point, if any of the cells has acquired a non-viable genotype as defined by our DAG or fitness landscape it is instantly removed from the simulation.
6. Finally, all alive cells have a chance to migrate to an empty space in it’s Von Neumann neighbourhood depending on the specified migration probability [34].

**Fig. 2.**
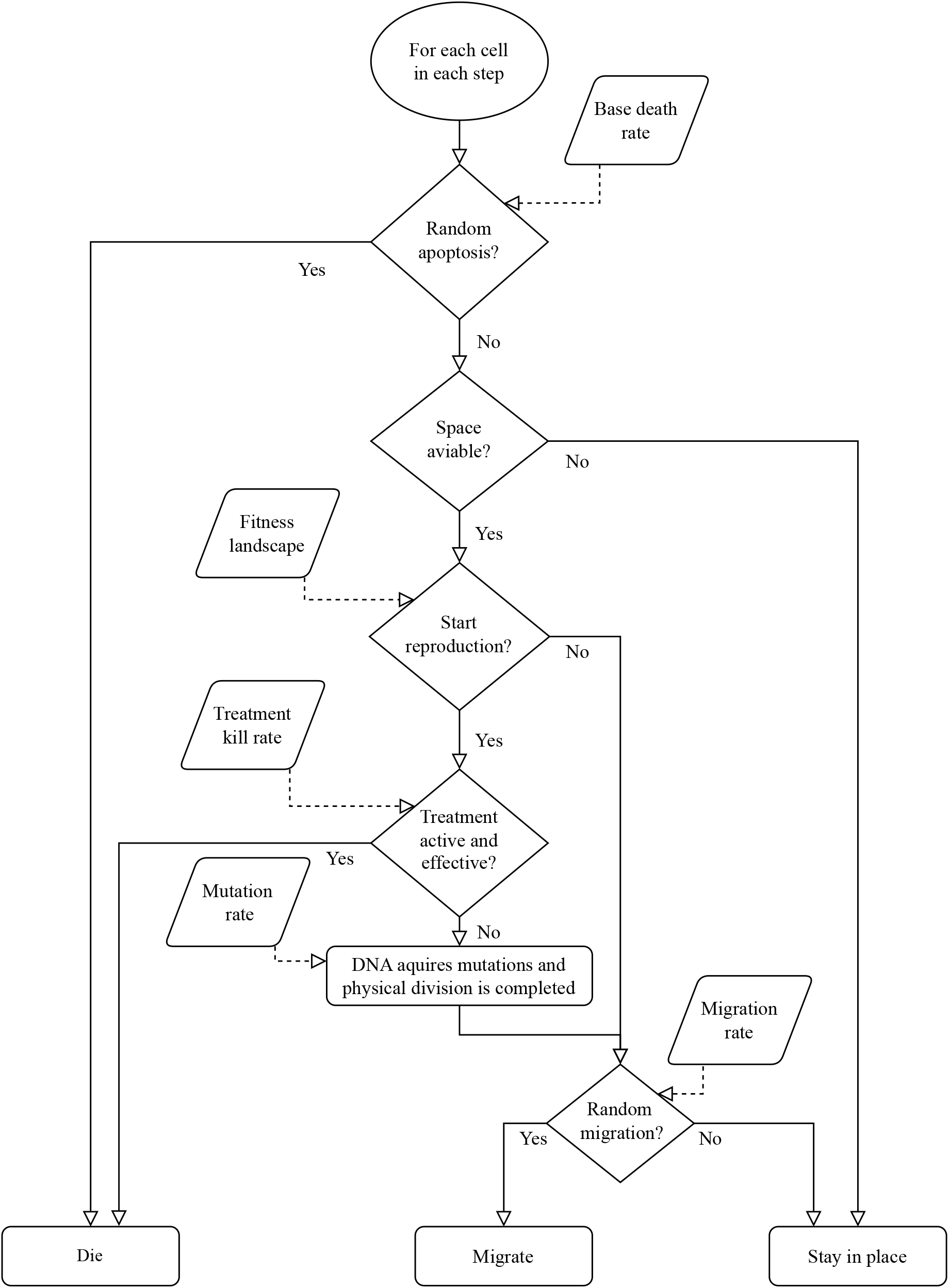
Decision making flowchart for each cell in each step. During each iteration of the ABM every cell is evaluated to decide whether it should die, reproduce or migrate according to the given model variables.

All of the steps of our model are designed to be consistent with other similar and already established spatial tumor development models found on the literature [38, 41, 46], allowing us to compare results between models and ensuring that all of the studied parameters can be interpreted in a similar way.

### 2.4 Adaptive therapy versus continuous therapy

The adaptive versus continuous therapy experiment shown in Figure 3 was based on the experiments performed by the group of Strobl et al. on 2022 [41]. We created a 2D simulation of a tumor where the starting conditions were 100 wild type cells in the center of a 200 by 200 lattice.

**Fig. 3.**
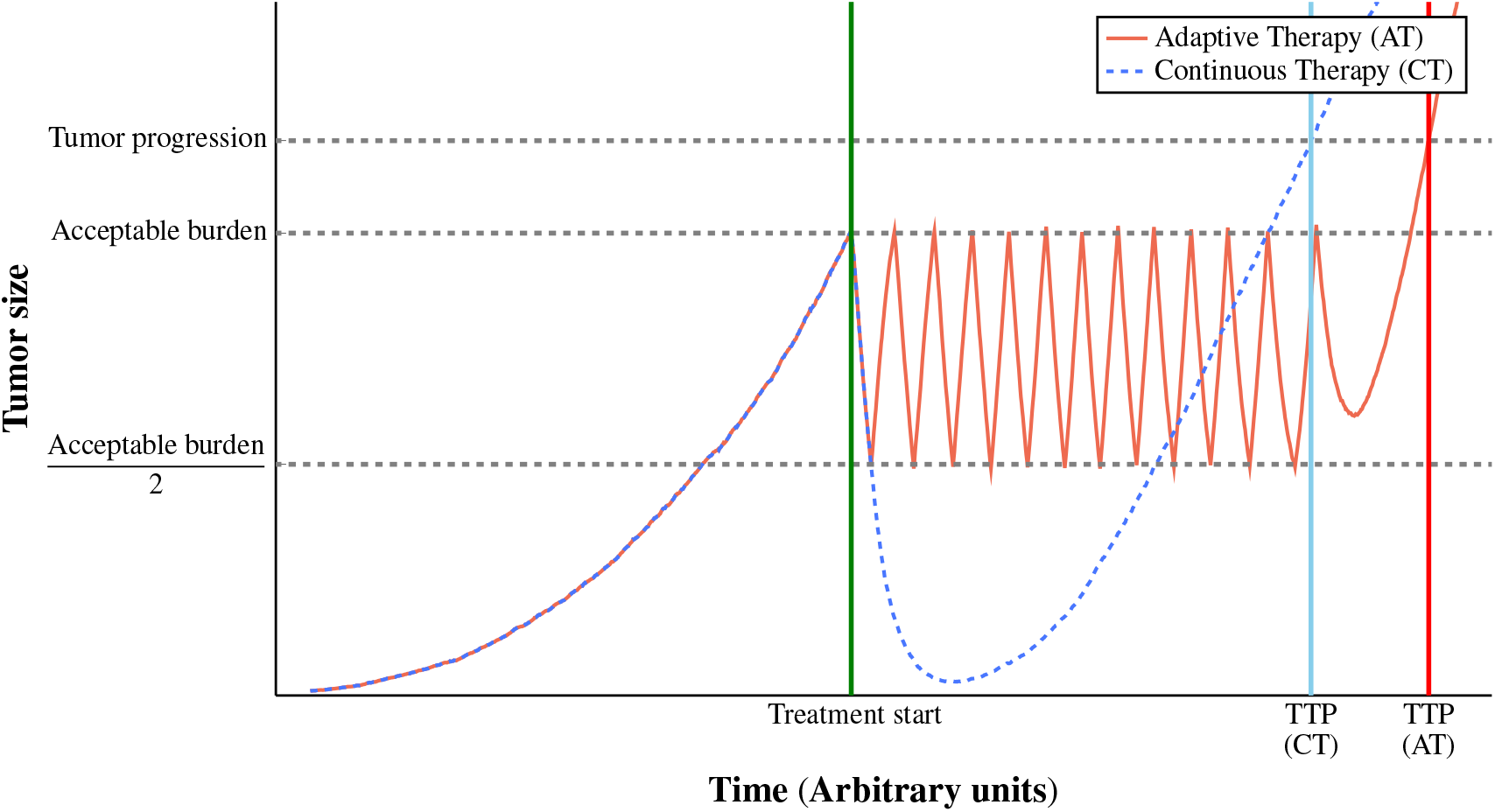
Performance of adaptive therapy versus continuous therapy. Time to progression two identical tumors, one treated under adaptive therapy versus other treated under continuous therapy. Treatment starts as soon as the tumor is detected and is paused at 0.5 times the acceptable burden size only in the case of adaptive therapy.

The values for the parameters of the simulation were chosen to be inside the biologically relevant ranges found in the literature [41] while also favouring adaptive therapy success. We chose a base death probability of 30%, a treatment kill probability of 75%, a mutation probability of 0.5% and a migration probability of 10%.

For the tumor treated under adaptive therapy we used the common treatment schedule of pausing the treatment at 50% of the original tumor size and letting it grow again until it reaches the acceptable burden size; when the acceptable burden size is reached, we start the cycle again [23, 40]. The schedule for the tumor treated under continuous therapy just consisted of a maximum continuous dose without pauses.

Both simulations are allowed to continue until they reach 1.2 times the original tumor size. At that size we consider we can no longer control tumor growth and we signal it as the TTP [40, 41].

### 2.5 Study of resistant cell’s growth inhibition

To study the genotypes inhibiting the growth of resistant cells, we ran 10.000 simulations for each of the studied DAGs or fitness landscapes, using the contact inhibition interaction rule. As values for the parameters of the simulations we used the same ones as for the adaptive versus continuous therapy experiment: 30% base death probability, 10% migration probability, 75% treatment kill rate and a treatment schedule pausing at 50% of the acceptable burden size.

For all of these experiments, the distribution of genotypes inhibiting the growth of resistant cells was calculated by normalizing the total sum of all genotypes that were present around resistant cells that failed the “Space aviable?” check of the algorithm shown in Figure 2.

As additional information to the figures showed in the results section, in Table 1 we show other relevant data from each of the scenarios studied in our simulations, like the mean treatment start and tumor progression times. We also include the mutation rate of each of each scenario, which was modified to adjust the “speed” of progression of each tumor to obtain a percentage of resistant cells on detection close to the 10^−2^ range. The percentage of resistant cells on detection has been shown in the literature to highly affect the outcome on adaptive therapy under spatial models [41], lowering the predicted increase in TTP as the percentage increases.

**Table 1:** Relevant data for each scenario used in the simulations of the resistant cell’s growth inhibition experiments. Treatment start is the moment the tumor reaches a large enough size to be detectable as defined in the treatment schedule. Tumor progression is defined as the moment at which we can no longer maintain the tumor under control (i.e. tumor size is 1.2 times over the acceptable burden). Resistant cells on detection is the percentage of cells that are resistant in the time step where the treatment starts.

| Scenario | Treatment start | Tumor Progression | Mutation rate | Resistant cells on detection |
| --- | --- | --- | --- | --- |
| Three genes with no restrictions | 504.6 | 840.2 | 0.0033 | 3,8% |
| Three genes with AND restrictions | 481.2 | 833.3 | 0.01 | 1,3% |
| Three genes with OR restrictions | 493.6 | 809.2 | 0.006 | 2,7% |
| Seven genes with no restrictions | 667.2 | 1095.3 | 0.005 | 3,6% |
| Seven genes with example 1 restrictions | 739.8 | 1115.8 | 0.02 | 3,4% |
| Seven genes with example 2 restrictions | 702.3 | 1169.4 | 0.01 | 1,6% |
| Seven genes with NK fitness landscape 1 | 587.7 | 920.9 | 0.01 | 2,8% |
| Seven genes with NK fitness landscape 2 | 721.2 | 1401.1 | 0.01 | 0,7% |

#### 2.5.1 Simulations with DAGs of three and seven genes

To transform the DAGs shown in Figure 4 into the table of mitosis probabilities for each genotype needed for our model we used the following approach:

**Fig. 4.**
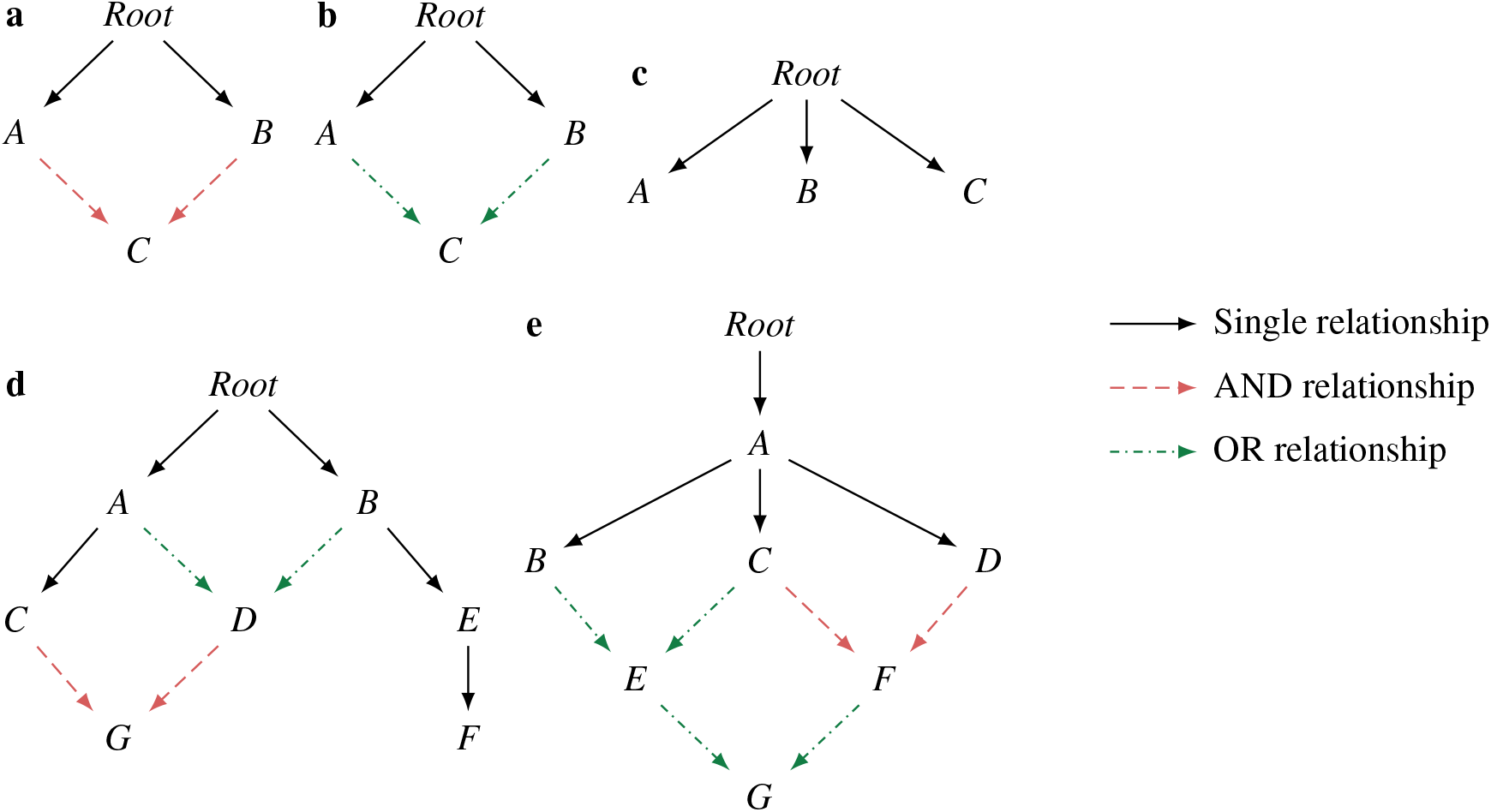
Restriction DAGs used in the simulations. **a**. Three genes with AND restrictions. **b**. Three genes with OR restrictions. **c**. Three genes with NO restrictions. **d**. Seven genes with example 1 restrictions. **e**. Seven genes with example 2 restrictions. See text for details about AND and OR restrictions. In the scenarios used here, in the three gene examples the resistant genotypes were those with gene C mutated, whereas in the seven gene examples the resistant genotypes were those with gene G mutated.

- We assumed the wild-type genotype had a mitosis probability of 2.7%, a value taken from other similar models [41].
- If a genotype was not allowed by the DAG, it had a mitosis probability of 0%.
- For all genotypes allowed, each non-resistant-conferring mutation had a multiplicative effect on the fitness, such that the genotype with all of the genes mutated except the resistant-conferring one, had a mitosis probability of 1.5 times that of the wild type.
- For all genotypes with the resistance-conferring mutation, a cost of resistance was applied, reducing the mitosis probability of the cell by 30%, a reasonable value found in the literature [40].

#### 2.5.2 Simulations with NK fitness landscapes

For the NK fitness landscape experiments, we used the OncosimulR [11] package to generate two random fitness landscapes under the NK model [29]. This model generates “tunably rugged” landscapes [47], allowing us to define N, the number of genes, and K, a measurement of the ruggedness of the landscape (the number of loci that interact with each locus). In our case we generated our fitness landscapes using the values of N=7 and K=3 to obtain a plausibly realistic result [12].

To avoid a trivial scenario where the resistant-confering mutation would be immediately accessible, from all of the landscapes we generated we decided to conduct the simulations using two landscapes where the fitness of the genotype that only had the resistance-conferring mutation “G” was significantly inferior to wild-type.

We then used the inbuilt functionality of the OncoSimulR package to convert the values of the fitness landscape to mitosis probabilities inside the range of 0% (no mitosis) and 4% (maximum mitosis), fixing the WT mitosis probability at a 2.7%, like in previous experiments.

#### 2.5.3 Data treatment: Jensen-Shannon divergence, scaling and plotting

To quantify the divergence between the relative frequencies of each genotype in the population and the real frequency of growth-inhibiting genotypes, we decided to use the Jensen-Shannon (JS) divergence, which can be defined as

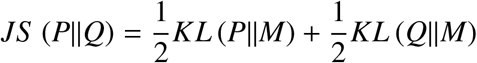

where 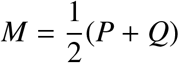 and KL is the Kullback–Leibler divergence.

The JS divergence is a symmetrized and smoothed version of the Kullback–Leibler (KL) divergence [8, 30]. It’s main advantage over the KL divergence is that we are still able to compute a value even in extreme cases where we might only have one genotype in the simulation, or where only one genotype is inhibiting the growth of resistant cells.

To obtain the results of Figure 6, Figure 7 and Figure 8, we then scaled the time axes using the data from Table 1 such that

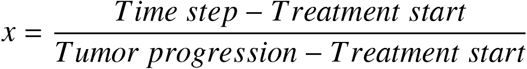

This normalization allows us to represent together simulations with different total durations and makes it easy to compare how the JS divergence evolves during the only timeframe where are actually able to make decisions, that is between the start of the treatment and the eventual tumor progression.

## 3 Results

### 3.1 Adaptive therapy versus continuous therapy

To test if our model agent-based model (ABM) was able to capture the benefits of adaptive therapy, we decided to recreate a similar experiment as the one conducted by Strobl et al. in 2022 [41]. In this simple experiment, we compared the TTP of two identical tumors, one treated with continuous therapy and one with adaptive therapy, using the common strategy of pausing the treatment once we reached half of the acceptable burden size.

As we can see in Figure 3, the tumor size is identical in both scenarios up until we start the treatment. When treatment starts, the tumor treated with continuous therapy is shrinked as fast as possible until the eventual appearance of resistant cells allows it to grow again. Meanwhile, in the tumor treated with adaptive therapy we maintain the tumor size contained between a lower and upper size limit, up until it also ends up acquiring resistance to the treatment. As we can see, the tumor treated with adaptive therapy achieves a longer TTP than the one treated with continuous therapy, which lines up with the results of other groups [19, 24, 40, 41] and supports the initial hypothesis in favour of adaptive therapy.

### 3.2 Study of resistant cell’s growth inhibition

After establishing that we were able to capture at least part of the phenomena responsible for the success of adaptive therapy, we decided to test whether restrictions in the order of accumulation of mutations had any effect on the results of the simulations.

We used a simple DAG model with “Single”, “AND” and “OR” dependencies to represent the restrictions needed to acquire a certain mutation. As an example, the model in Figure 4a can be interpreted as follows: We can see that mutations A and B have a “Single” dependency on the “Root” node, meaning they can be acquired at any time independently of each other. However, mutation C has an “AND” dependency on mutations A and B, meaning that it can only be acquired after both mutations A and B are already present. If we instead take a look at Figure 4b, we can see that mutation C has an “OR” dependency on mutations A and B, meaning that it just needs either A or B mutations present before it can be aquired. All of the DAGs used in the simulations can be seen in Figure 4. CPM methods that fit models with AND restrictions include CBN [20], OncoBN [33] and PMCE/H-ESBCN [2]; CPM methods that fit models with OR restrictions include OncoBN and PMCE/H-ESBCN; further examples of these models, and discussion about generating fitness landscapes from them, are given in the supplementary information of EvAM-Tools [13].

In Figure 5 we can see the results of a single simulation ran using three genes with the AND restrictions DAG shown in Figure 4a. In this simulation, the genotype with gene C mutated is the resistant genotype. Figure 5b represents the relative frequency of each genotype. We can see that at the start of the treatment we have an heterogeneous tumor with multiple cell populations coexisting, including a small percentage of resistant cells. This percentage of resistant cells rises as we continue treating the tumor until it becomes the predominant population and we lose control of the tumor size.

**Fig. 5.**
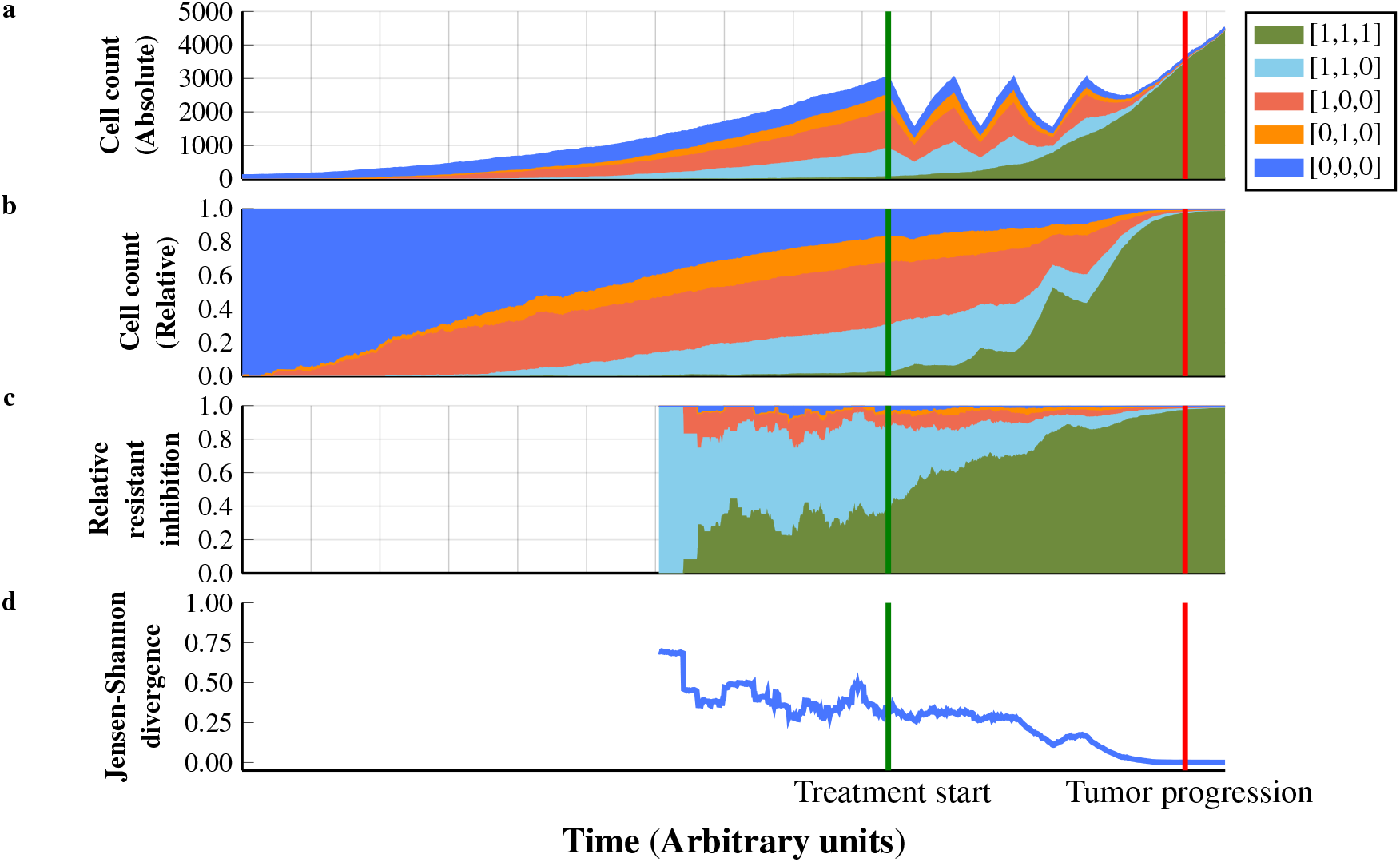
Deviation of real competition from perfect mixing model under the three genes with AND restrictions DAG. Genotypes are represented as [A,B,C] where 0=WT and 1=Mutated. For example, genotype [0,1,0] is the genotype with only gene B mutated. The resistant genotype is the genotype with gene C mutated, [1,1,1]. **a**. Absolute frequency of each genotype in the simulation. **b**. Relative frequency of each genotype in the simulation. **c**. Relative source of growth inhibition on resistant cells, as defined in section 3.2. **d**. Jensen-Shannon divergence between the relative frequency of each genotype shown on **b** and the relative source of growth inhibition of resistant cells shown in **c**.

**Fig. 6.**
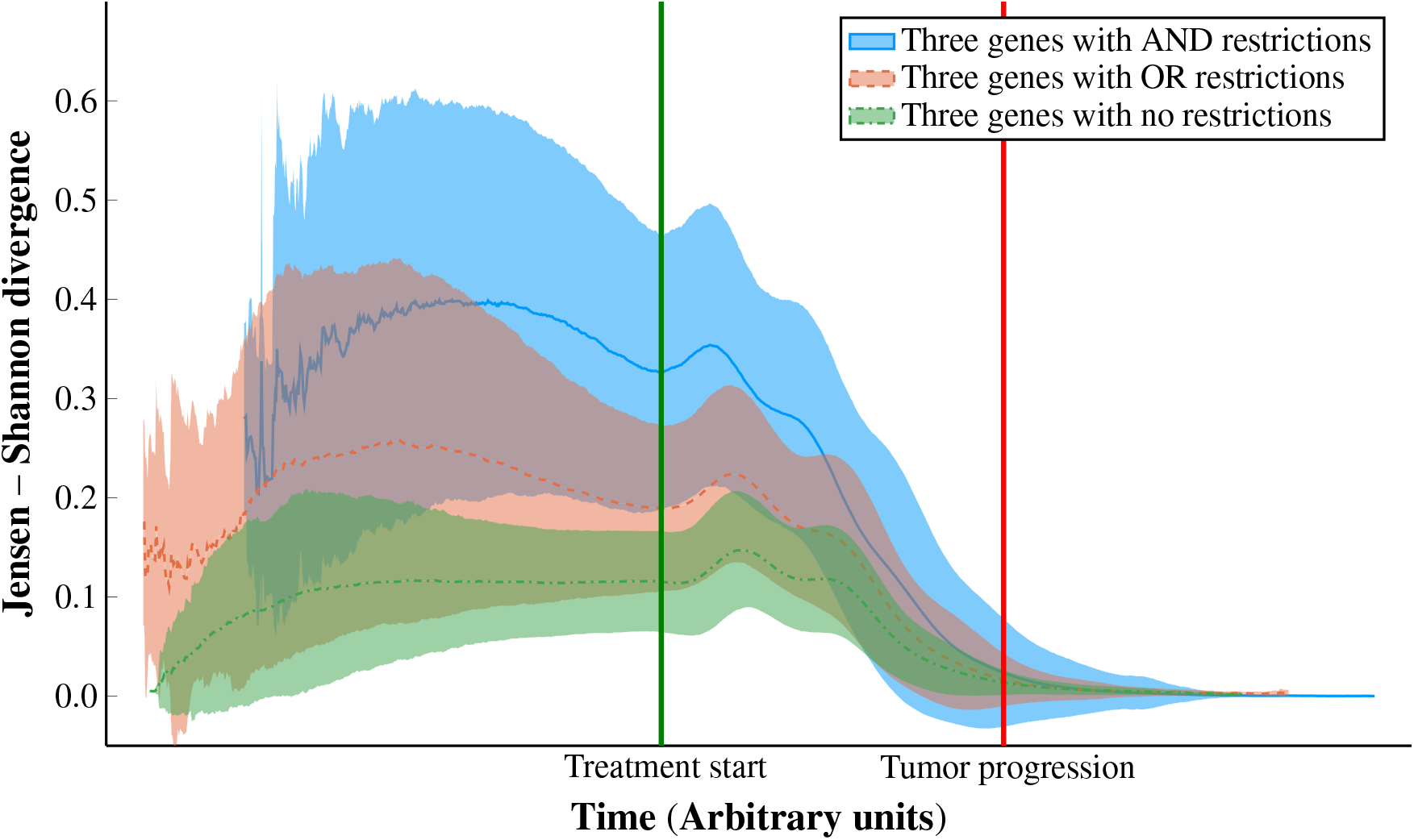
Deviation of real competition from perfect mixing model using DAGs of three genes. Jensen-Shannon divergence between the relative frequency of cells in the simulation (perfect mixing model) and the real relative source of inhibition. Figures are not lined up on the left because of the scaling of the time axis —see section 2.5.3— combined with differences in time of Treatment start, as shown in Table 1.

**Fig. 7.**
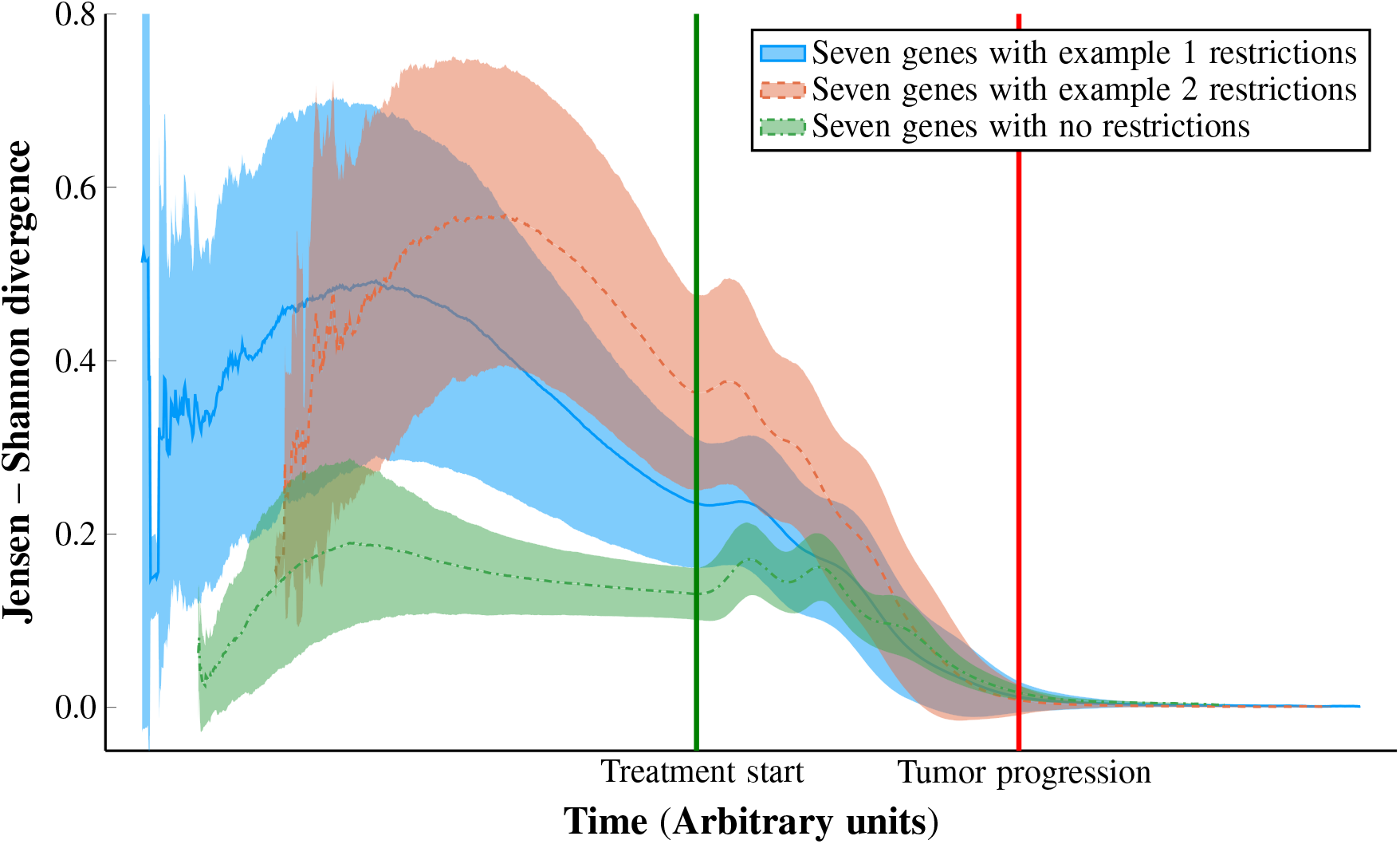
Deviation of real competition from perfect mixing model using DAGs of seven genes. Jensen-Shannon divergence between the relative frequency of cells in the simulation (perfect mixing model) and the real relative source of inhibition. See Fig. 6 for additional details.

**Fig. 8.**
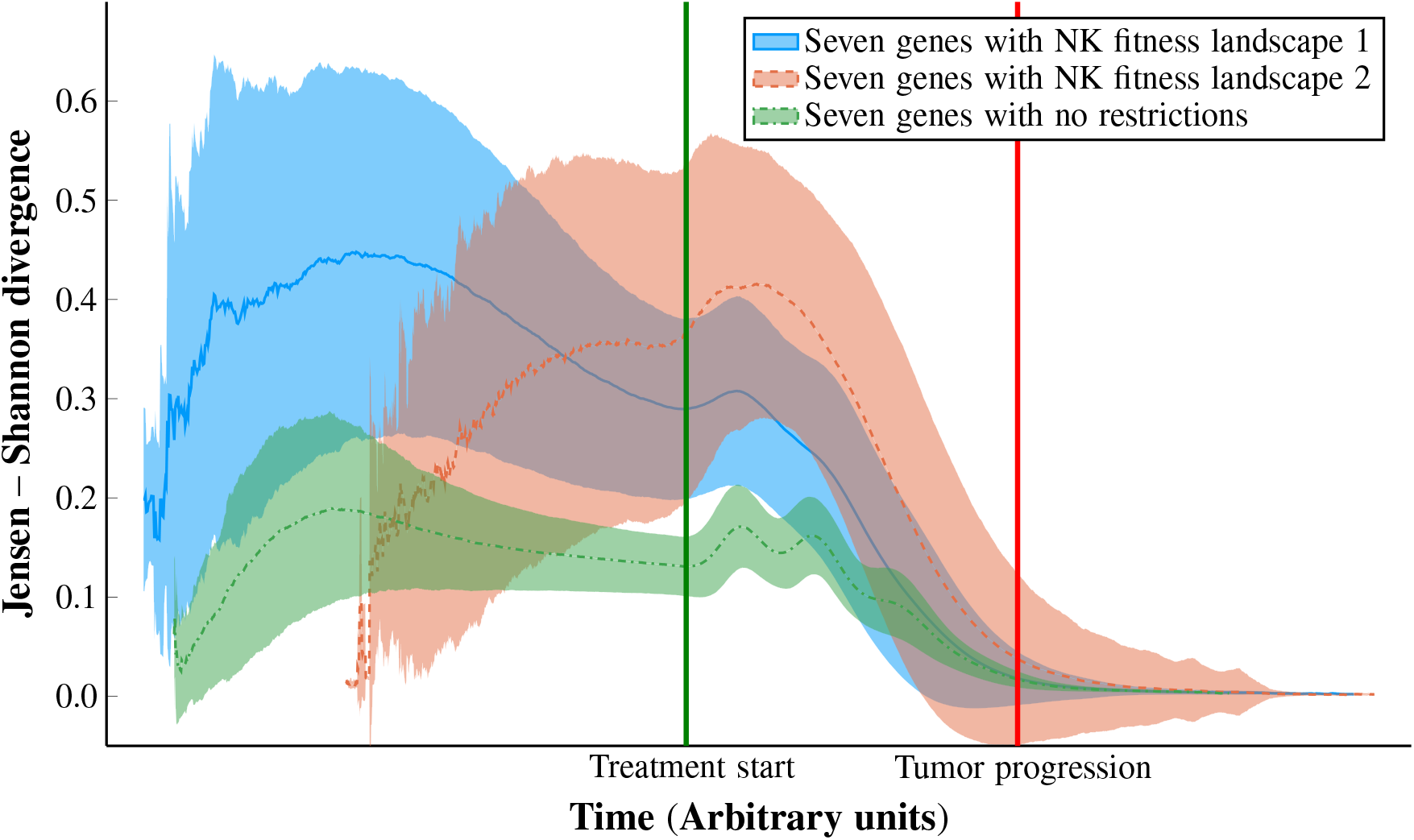
Deviation of real competition from perfect mixing model using two fitness landscapes generated under the NK model. Jensen-Shannon divergence between the relative frequency of cells in the simulation (perfect mixing model) and the real relative source of inhibition. See Fig. 6 for additional details.

Interestingly, the distribution of genotypes responsible for the growth inhibition of the resistant cells, as shown in Figure 5c, is significantly different to the relative frequency of each genotype on the population, suggesting that the genotype of the cells inhibiting the growth of the resistant cells is not random, and is heavily influenced by the restrictions in the order of accumulation of mutations.

To numerically quantify this discrepancy between the relative frequency of the genotypes and the actual inhibition from each genotype, we calculated the Jensen-Shannon divergence [8, 30] between the two distributions. In our specific case, we can interpret this metric as how different the distribution of cells inhibiting the resistant cells is from what we would expect from a model that uses the perfect mixing assumption. The results for this simulation can be seen in Figure 5d.

#### 3.2.1 Simulations with DAGs of three genes

As a first approach to study the discrepancy between the relative frequency of the genotypes and the actual inhibition from each genotype, we constructed three simple restriction DAGs with three genes each, which are shown in Figure 4, and established the mutation in gene “C” to be the one that confers resistance to our treatment.

As we can see in Figure 6, as we impose harder restrictions on the accumulation of mutations (No restrictions < OR restrictions < AND restrictions), we get a larger deviation of the real competition from the one we would expect in a perfectly mixed model.

The use of this very simple model with three genes allows us to reason about the results shown in Figure 6. In the case of the DAG with AND restrictions, the resistance mutation can only appear in zones of the tumor where A and B are already mutated, making that genotype it’s main competitor even if it is actually relatively infrequent in the population. In the case of the DAG with OR restrictions, the resistance mutation will only appear in zones where either A or B are mutated, making it a softer restriction and thus more similar to the perfect mixing assumption. Finally, in the DAG with no restrictions, where the resistance mutation is always accessible, the distribution is very similar to that of a perfectly mixed model.

#### 3.2.2 Simulations with DAGs of seven genes

Although the simulations with three genes allows us to easily interpret the results, they can be excessively simplistic compared to the real world. To examine whether our results would hold up in more complex scenarios, we created two DAGs with seven genes each, represented in Figure 4. Each of these DAGs contains simultaneously “Single”, “AND” and “OR” relationships, making the possible mutational paths that lead to the resistance mutation G much more diverse than in the previous experiment.

As we can see in Figure 7, even in this more complex scenario we can still see that the distribution of genotypes inhibiting the growth of resistant cells is not random, and is dependent on what restrictions are present. Unfortunately, as we increase the number of genes on our model the number of genotypes grows up exponentially, reaching 128 possible genotypes at seven genes (128=2^7^). This means that it is no longer trivial to explain why a given DAG has a higher deviation compared to another one.

#### 3.2.3 Simulations with NK fitness landscapes

Finally, we wanted to examine if we could still observe this discrepancy between the relative frequency of the genotypes and the actual inhibition from each genotype under more realistic fitness landscapes, confirming it was not an artifact of our DAG restriction model. To examine this we generated two random NK fitness landscapes [29] of seven genes using the OncoSimulR package [11]. In these simulations, the resistant genotypes were those with the gene G mutated.

The results shown in Figure 8 confirm that there is still a significant difference in the genotypes inhibiting the growth of the resistant clones compared to a perfectly mixed model, at least in a similar magnitude as the ones seen in the experiments with the more simplistic DAG model. However, the results obtained with the NK fitness landscapes are the least interpretable out of the three experiments and probably the most difficult to extract conclusions from.

## 4 Discussion

### 4.1 Study of resistant cell’s growth inhibition

A large number of state-of-the-art models [48] assume that all the cells in the tumor are perfectly mixed, as it allows us to develop simpler mathematical models and borrow tools from fields like ecology such as the Lotka–Volterra equations. Under this assumption, the distribution of genotypes interacting with any given cell is exactly the relative frequency of the genotypes in the population. However, this is an oversimplification that may be specially misleading when modelling adaptive therapy, as knowing the phenotype of the cells the resistant cells are competing with can be vital when predicting the success or failure of this approach [40, 45].

It has recently been pointed out that often, the most common genotype inhibiting a resistant cell is the genotype of the resistant cell itself [41]. This is expected, as when reproducing, daughter cells usually remain close in space after mitosis is completed. In our work, we expand on this remark, and show that when we take into account the spatial architecture of the tumor and restrictions in the order of accumulation of mutations, the difference between the perfect mixing assumption and reality can be higher than previously anticipated, even when only considering the interaction with sensitive cells.

One of the highest risks of not taking into account the discrepancy between the relative frequency of the genotypes and the actual inhibition from each genotype can manifest itself when making predictions on real patients. Most of the proposed protocols for predicting adaptive therapy success [40] obtain the measurements for the properties of the tumor either by using an homogenate of the whole tumor or by taking the mean of results from different parts of the tumor, which is in essence a weighted average of the specific properties of each genotype (fitness, base death rate…) by the relative frequency of each of those genotypes in the whole tumor. As we have shown, if we assume that these properties are representative of those of the cells interacting with resistant cells, we can be making highly inaccurate predictions, specially in tumors with high heterogeneity.

Our simulations, as presented in Figures 6, 7 and 8, show that this divergence is significantly higher at the start of the treatment and decreases as we start treating the patient and the resistant genotypes start taking over the tumor. This means that this information can be used during the open window of action between the detection of the tumor and the eventual progression. This is important specially when viewing adaptive therapy from the lens of game theory, where the oncologist plays as a rational agent and cancer cells are only able to adapt to the situation created by the oncologist [39]. In this scenario, every bit of information can give the oncologist an advantage that could improve the prognosis of the patient.

Our understanding is that this effect has not been studied before in the literature. Thus, more work is needed to understand how it affects already developed models of adaptive therapy. Specially, more work is needed to understand how we can use this newly found phenomena to improve our predictions on adaptive therapy success and to create better treatment schedules and improve TTP. We think the use of CPM models might prove particularly useful in this field as their ability to infer restrictions in the order of mutations can serve both as a predictor of the risk of acquiring resistance mutations, and as a tool to predict the genotypes of cells interacting with these resistant cells.

### 4.2 Additional possible use cases of our model

Apart from the data used in the experiments shown in this work, our model is able to output other potentially useful data like cell phylogeny or population diversity. Additionally, the model has been built in a highly modular way, allowing us or other researchers to easily implement any other feature we might need for future experiments easily and without interfering with existing functionalities. All of this makes our model a viable tool for studying adaptive therapy in all scenarios where there might be an interest in studying the spatial architecture of the tumor, such as in the case of restrictions in the order of accumulation of mutations.

Some of the possible research use cases for our model are the following.

- Investigate how the spatial architecture of the tumor, including the location of the different sub-populations, can affect tumor development and prospects of survival, either under continuous or adaptive therapy.
- Test how different fitness landscapes can affect tumor development under different modes of tumor evolution [34], and their effect on the predicted TTP of the tumor.
- Check if the already studied parameters for predicting adaptive therapy success, like cell turnover [40] or cost of resistance [19], are still able to make good predictions when we introduce a spatially explicit model or different fitness landscapes.
- Use the data from the prediction of genotypes inhibiting the growth of resistant cells to develop new algorithms able to make better informed decisions.
- Study the effect of multi-drug treatment [23], drug dose modulation [18] or changes in the acceptable burden size [23] to help design better treatment schedules that are able to better control the tumor and prolong the life of the patient.
- Investigate the risk associated with maintaining a high number of sensitive cells that are themselves able to become resistant during the course of adaptive therapy, an effect that can actually result in a harmful outcome for the patient and a decrease in TTP [24].

Apart from the traditional research that can be conducted with this model, we also propose it as a potential teaching tool that will allow both students and researchers to familiarize themselves with concepts such as tumor heterogeneity and architecture, fitness landscapes and adaptive therapy. Our model allows us to run simulations in real time and change the parameters of the simulation on-the-fly, letting the user experiment with different parameters and gain intuition on how they may effect the development of the tumor. Right now the model supports rendering simulations both in 2D and in 3D, as can be seen in Figure 9.

**Fig. 9.**
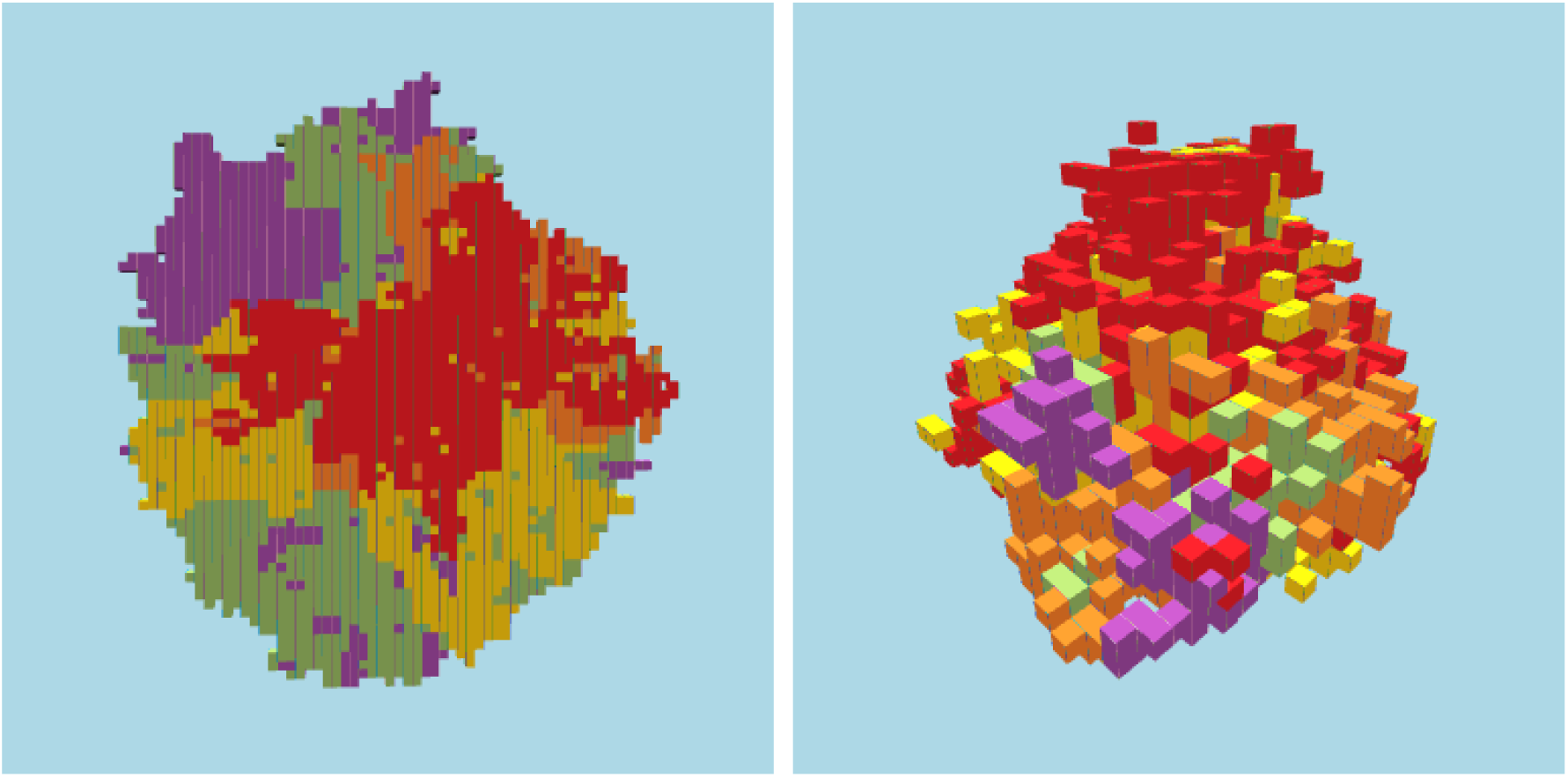
Snapshots of 2D and 3D tumors simulated with our agent-based model. Both images were taken just before the treatment starts and correspond to tumors simulated with the contact interaction rule and the three genes with AND restrictions DAG. Each color represents a different genotype.

Our model, however, also has its limitations. As mentioned earlier, agent-based models often require powerful hardware to run a high number of simulations in an acceptable time frame. Additionally, our model makes some assumptions that may make it unable to capture some of the phenomena observed in real tumors. One of those assumptions is that changes in the treatment take effect instantaneously and that the concentration of the treatment is constant across the tumor. Furthermore, our model does not explicitly take into account other spatial properties of the tumor that may impact the spatial architecture such as vasculature, cell signaling or nutrient consumption. These properties, however, could be added to the model if needed thanks to the modularity of the code.

## 5 Conclusion

We have developed a spatially explicit agent-based model to examine the effects of restrictions in the order of accumulation of mutations on adaptive therapy. With this model, which can also be used as a teaching and research tool to address other questions about adaptive therapy and evolutionary dynamics, we have shown that restrictions in the order of accumulation of mutations can have a large effect on the competitive environment that resistant cells experience. Our work emphasizes the need to consider these restrictions when designing adaptive therapy protocols and highlights the need for combining these two areas in further modeling and simulation work.

## 6 Code availability

Code and scripts available from https://github.com/YM162/TumorSim under the GNU GPL v. 3.0 license.

## 7 Acknowledgements

Supported by grant PID2019-111256RB-I00 funded by MCIN/AEI/10.13039/501100011033 to Ramon Diaz-Uriarte.

